# Suboptimal refeeding compensates stunting in a mouse model of juvenile malnutrition

**DOI:** 10.1101/2024.03.25.586077

**Authors:** Jean-Louis Thoumas, Amandine Cavaroc, Damien Sery, François Leulier, Filipe De Vadder

## Abstract

**Background:** Early life, particularly after weaning, is the most rapid period of growth in mammals, and this growth is highly dependent on adequate nutrition. Protein-energy malnutrition (PEM) during this critical window can lead to stunting and wasting, which have long-term health consequences.

**Objective:** This study aimed to develop a mouse model of juvenile PEM to assess the effects of refeeding with various diets and interventions on growth recovery, including the impact of probiotic supplementation and suboptimal refeeding diets.

**Methods:** Juvenile C57Bl/6J mice were fed a low-protein diet (LPD, 5% kcal from protein) starting at postnatal day 14 (P14) to induce malnutrition. Following weaning, both male and female mice were refed an optimal diet (Altromin 1310, 27% kcal from protein) at different times ranging from P28 to P56. In a second intervention, male mice were supplemented during refeeding with *Lactiplantibacillus plantarum* WJL (Lp^WJL^), a probiotic known to stimulate growth in malnourished conditions. A final group of malnourished male mice were refed with a Western diet (WD, 34.5% kcal from fat; 15.3% kcal from protein) or a modified Western diet (MWD, 34.2% kcal from fat; 7.5% kcal from protein) to model suboptimal refeeding.

**Results:** Refeeding with an optimal diet fully restored growth in female mice, but male mice exhibited persistent stunting despite nutritional rehabilitation. Lp^WJL^ treatment during refeeding did not enhance systemic growth in males. In contrast, refeeding with WD or MWD restored body length but impaired glucose metabolism, particularly in mice refed MWD after PEM. Lp^WJL^ exacerbated glucose intolerance in the suboptimal refeeding groups.

**Conclusion:** Sex-dependent differences exist in the recovery from early-life malnutrition, with males showing incomplete growth recovery despite optimal refeeding. Suboptimal diets, while compensating for stunting, impair glucose metabolism, especially when protein intake is insufficient. Probiotic supplementation with Lp^WJL^ did not improve growth outcomes.

## Introduction

Malnutrition in the early postnatal period is a significant concern in many developing countries and is regarded as the leading cause of death among children under the age of five (1). Notable outcomes include wasting and stunting, defined by a child’s weight-for-height and height-for-age being more than two standard deviations below the median according to the World Health Organization’s Child Growth Standards. Wasting is indicative of acute growth disturbance from malnutrition, whereas stunting is more symptomatic of chronic malnutrition, with impaired long-term growth (2). In 2022, about 148 million children under five years of age were stunted (1). Stunting is directly caused by an inadequate intake of nutrients, especially dietary protein (3). Besides stunting, severe malnutrition in early life is associated with metabolic disturbances that persist later in life (4,5).

Extensive epidemiological studies have demonstrated that malnutrition during early life has detrimental effects on health, which can last throughout the life of the malnourished person (6). This is called nutritional or metabolic programming, a process through which metabolic and physiological adaptations in response to a poor diet persist even after the nutritional stress is removed (7–10). The first 1,000 days of life are crucial for growth because of the high developmental plasticity during that time, after which stunting generally becomes irreversible and leads to growth impairment in adulthood (11,12). Recently, three articles of large human cohorts have highlighted the importance of early life interventions to prevent wasting and stunting in more than 30 cohorts of malnourished children (13–15).

In this context, establishing animal models of early life malnutrition seems crucial to understand the mechanisms behind the observed adverse consequences. Previously, we established a model of mouse juvenile malnutrition, upon which mice were fed a low-protein low-fat diet from weaning (post-natal day 21, P21) to eight weeks of age (16,17), which is the rough physiological equivalent of 1 year to 20 years of age in humans (18). We showed that supplementation with the probiotic *Lactiplantibacillus plantarum* WJL (Lp^WJL^) sustains postnatal growth of malnourished animals by stimulating anabolic metabolism and hormonal changes, namely the somatotropic axis via the secretion of insulin-like growth factor 1 (IGF-1) (16,17). However, knowledge on the long-term consequences of early malnutrition after refeeding is lacking. Indeed, the first intervention to stop malnutrition is resolving the nutritional stress associated to the lack of protein by providing an optimized diet.

Using a mouse model of juvenile protein energy malnutrition, we sought to investigate the consequences on growth catch-up after refeeding with either an optimal or suboptimal diets (richer in fat and poorer in protein). While growth was not affected by a suboptimal diet, we also explored the metabolic alterations associated to this switch.

## METHODS

### Animals

Male and female specific pathogen-free C57Bl/6J mice, aged 9 to 10 weeks, were purchased from Charles River (L’Arbresle, France). After one week of acclimatization at the conventional animal room, they were mated in our animal facility and offspring pups were used for experiments. The number of mice per cage ranged from 2 to 6, depending on the sex ratio and the number of offspring. Mice were kept in Innorack IVC Mouse 3.5 disposable cages (Inovive, USA), exposed to 12:12 hours light-dark cycles, supplied with tap water and given free access to food. The mice’s environment was enriched with paper towels for nesting, Aspen blocks, mouse densified cellulose domes (Genobios, Laval, France), as well as Innorichment enrichment sheets (NCE LifeSciences, Ahmedabad, India). These items promote the mice’s well-being and natural behaviors. All experiments were performed in accordance with the European Community Council Directive of September 22, 2010 (2010/63/EU) regarding the protection of animals used for experimental and other scientific purposes. The research project was approved by a local animal care and user committee (C2EA015) and subsequently authorized by the French Ministry of Research (APAFIS#22788-2019091713204925 v8 and APAFIS#32380-2021061809124474 v4).

### Experimental design

Specific-pathogen-free C57Bl/6J mice were mated. After delivery, the litter size was reduced to six offspring per dam. Experimental groups were formed based on litter size and the timing of pup birth, with no randomization applied. All experimenters were aware of the experimental groups throughout the study. From birth to post-natal day 14 (P14), the dams were given free access to a breeding diet (optimal diet; Altromin 1310). At P14, the pups were kept with their mother, but the food was changed according to the experimental groups. All diets were irradiated at 40 kGy before use. Composition of the experimental diets is given in Table 1. At P21, mice were weaned on the diet introduced at P14. Experimental sample size was determined in order to reach a statistical power of at least 80%, using G*Power 3.1 (19). Body weight and length were determined once a week. For the measurement of body length, mice were briefly anesthetized with isoflurane and length was measured using a graduated ruler. Body length was defined as the nose-to-anus distance. Starting at day 21, we estimated food intake by measuring the initial amount of food provided and the remaining food after one week, then calculating the average intake per mouse in each cage. At the end of the experiments, mice were killed by cervical dislocation. Tissues were sampled (Figure 1D), weighed and measured, and snap-frozen in liquid nitrogen.

**Figure 1:**
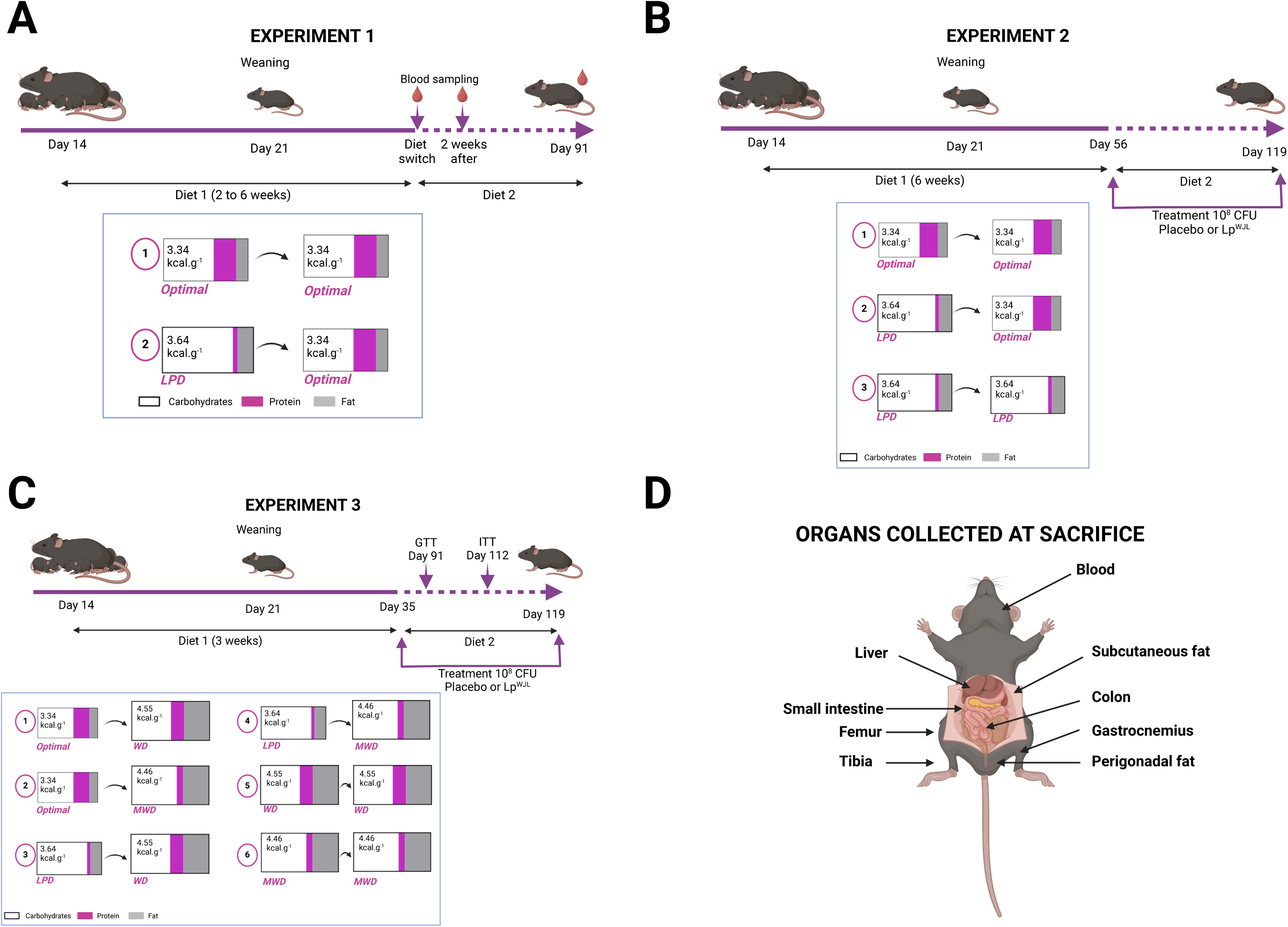
Overview of experimental procedures. **(A)** *Experiment 1*: Mice were fed a low-protein diet until P28, P35, P42, P49, or P56, then refed an optimal diet until P91, with blood samples collected at key time points. (**B**) *Experiment 2*: Optimal, low-protein, and refeeding groups were given placebo or *Lactiplantibacillus plantarum* WJL (Lp^WJL^) from P56 to P119. (**C**) *Experiment 3*: Mice were fed an optimal, low-protein (LPD), Western (WD), or modified Western diet (MWD), and treated with placebo or Lp^WJL^ from P35 to P119. (**D**) Organs were collected at sacrifice for further analysis.

Experimental procedures are summed up in Figure 1.

#### Experiment 1 (male and female mice) – *Figure 1A*

A group of mice was given optimal diet (Altromin 1310) from P14 to P91. Experimental groups were given low-protein diet (LPD, composition in Table 1), which is a modified isocaloric version of AIN-93G diet. LPD was custom-made by Sciences de l’Animal et de l’Aliment de Jouy unit (SAAJ, INRAE UE1298, Jouy-en-Josas, France). Five groups of mice were switched from a low-protein diet to an optimal diet at different time points—P28, P35, P42, P49, and P56—and then maintained on the optimal diet until P91. Blood samples were taken from the submandibular vein three times: at diet switch (P28, P35, P42, P49, or P56), two weeks after the switch (P42, P49, P56, P63, or P70), and at P91. Time-matched samples were taken in the optimal-fed group. In this experiment, N = 8-10 mice per refeeding group. Each mouse was only bled three times throughout life. Thus, in order to have time-matched samples for optimal-fed mice, the optimal-fed group contained 25 males and 38 females.

#### Experiment 2 (males only) – *Figure 1B*

Three groups were formed: a group fed optimal diet until P119; a refeeding group, fed LPD from P14 to P56, then optimal diet from P56 to P119; and a low-protein group fed LPD from P14 to P119. From P56 to P119, two treatment arms were formed in each group: one receiving a placebo solution (maltodextrin) and one receiving 10^8^ CFU *Lactiplantibacillus plantarum* WJL (Lp^WJL^) dissolved in maltodextrin solution, as described previously (17). A 100 µL dose of the experimental solutions was administered orally five times a week in the morning. The solutions were delivered using a pipette, allowing the mice to lick the pipette tip. At P112, mice were taken to ANIPHY Platform (Lyon) and put in the Minispec LF90 II (Bruker, Wissembourg, France) for non-invasive determination of body composition using NMR. Mice were taken back to our animal facility and given the experimental diet until P119. In this experiment, N = 7-10 mice per group.

#### Experiment 3 (males only) – *Figure 1C*

A group was fed optimal diet from P14 to P119. Experimental diets were optimal diet (Altromin 1310), LPD, Total New Western Diet (WD; Teklad TD.140203) and a modified isocaloric version of WD (MWD), with reduced protein content. MWD was custom-made for the experiment. WD and MWD were purchased from Envigo (Gannat, France). WD is based on Total Western Diet, as described in (20), with the modifications described by Monsanto et al. (21). Composition of the diets is given in Table 1. As in Experiment 2, mice were given either a placebo or an Lp^WJL^ solution, from P35 to P119. In this experiment, N = 7-11 mice per group.

### Glucose and insulin tolerance tests

In Experiment 3, at P91 and P112, mice underwent respectively a glucose (GTT) or insulin tolerance test (ITT). Mice were food-deprived for 6 hours before the start of the tests. For the GTT, mice were given an intraperitoneal (i.p.) injection of a 20% sterile glucose solution (1 g.kg^-1^ body weight). For the ITT, mice were given an i.p. injection of human insulin (Novorapid, Novo Nordisk) at 0.75 U.kg^-1^ body weight. Glycemia was monitored at the tip of the tail vein at times 0, 15, 30, 45, 60, 90, and 120 minutes post-injection using a glucometer (One Touch Verio Reflect, Life Scan).

### Ethical limitations and endpoints

The following endpoints were established to ensure animal welfare during the study: *Severe Weight Loss*: Animals losing 20% or more of their maximum body weight during the experiment were removed from the study.

#### Behavioral Changes

Animals exhibiting reduced locomotion or lack of response to external stimuli were considered to have reached an endpoint.

#### Degraded Physical Appearance

Animals with severely deteriorated external physical condition, including rough fur and a hunched posture, were removed.

#### Excessive Weight Gain

For animals on a high-fat diet, those gaining more than twice the weight of age-matched control animals on a standard diet (as determined in prior experiments) were excluded.

#### Monitoring Schedule

A total of 322 mice were included in the study.

From postnatal day 7 (P7) to day 21 (P21), animals were weighed weekly.

From P21 until the end of the study at P119, weight and body length were measured weekly.

Daily general health observations were conducted throughout the study.

#### Insulin Tolerance Test (ITT)

During the ITT, any animal presenting a blood glucose level ≤1.67 mM (30 mg/dL) was immediately injected with glucose (150 μL of a 25% (w/v) solution) and temporarily withdrawn from the experiment. If the animal’s blood glucose and behavior normalized within 6 hours, it was reintegrated into the study. According to these criteria, 11 animals were temporarily excluded but were successfully reintegrated after recovering normal glycemia and behavior.

#### Glucose Tolerance Test (GTT)

During the GTT, animals with blood glucose levels ≥16.7 mM (300 mg/dL) two hours after glucose injection were killed.

#### Outcomes

All other criteria were met for the remainder of the study population.

### IGF-1 measurement in serum

Blood samples were collected from the submandibular vein at the time of diet switch, two weeks after the switch and at the end of the experiment, into a sample tube containing a clotting activator (Micro sample tube serum gel, 1.1 mL, Sarstedt). After centrifugation at 10,000 g (Eppendorf 5424 R) for 5 minutes, sera were collected into sterile 1.5 mL microtubes and kept at −70°C for further use. IGF-1 in serum was measured using Mouse/Rat IGF-1 Quantikine ELISA (R&D Systems) according to manufacturer’s instructions.

### Data treatment and statistics

Statistical analysis was performed using GraphPad Prism software version 10.0 (GraphPad, San Diego, California, USA). For Experiment 1, we used one-way ANOVA followed by Tukey’s multiplicity-adjusted test. Growth catch-up slopes were compared using ANCOVA. Principal Component (PC) Analysis was performed based on the morphological features measured, with PC selection based on percent of total explained variance. For Experiment 2, we used two-way ANOVA followed by multiplicity-adjusted Šídák’s test. For Experiment 3, we used one-way ANOVA followed by Tukey’s multiplicity-adjusted test when comparing to optimal-fed and two-way ANOVA followed by multiplicity-adjusted Šídák’s test when comparing treated groups. Unless stated otherwise, results are presented as mean ± SEM. *P*-values are reported after adjustment for multiple comparison. A value of *P* < 0.05 was considered statistically significant.

## RESULTS

### Optimal refeeding after long-term juvenile malnutrition is insufficient to fully recover growth in males, but not in females

In order to model protein energy malnutrition (PEM), we designed an AIN-93G-based isocaloric low-protein diet (LPD), with 5.1% total kcal coming from protein (Table 1). We first investigated how relevant our model of malnutrition was to human disease. Indeed, although several models have been published (22,23), there is no standard murine model of PEM. We first characterized the impact on weight gain and linear growth in C57Bl/6J mice fed an LPD, starting at P14 and going on after weaning (which took place at P21). We established five distinct groups, each corresponding to a different duration on the low-protein diet, ranging from 2 to 6 weeks. From P28 to P56, mice were refed with optimal chow diet. We defined our baseline comparison group as mice fed a chow diet, identified as the optimal diet for promoting growth in juvenile mice, as shown in Supplementary Figure 1, where growth on chow was superior to growth on the AIN-93G diet. To parallel juvenile human malnutrition, we defined stunting and underweight in our model as body size and weight falling more than two standard deviations below the mean of the optimal-fed group (dashed lines, Figure 2A-D). As expected, LPD-fed mice had severe growth retardation following weaning, associated to reduced body weight (Figure 2A-D). Since definition of wasting (low weight-for-length) in humans is complex and requires a Box-Cox power transformation (24), and growth does not occur in the same manner in mice and humans, we plotted length vs. weight in our experimental animals. We found that weight and length are linearly correlated during growth (Figure 2E-H). In LPD-fed mice, the slope was significantly smaller than in age-matched optimal-fed mice (males and females: ANCOVA: P < 0.0001), meaning that the mice were wasted (Figure 2E-F). In females, when mice were refed an optimal chow diet after LPD, growth and weight catch-up occurred within three weeks of diet switch (Figure 2B, D, H). However, in males, the longer the mice remained on LPD, the slower growth and weight catch-up occurred (Figure 2A, C). Indeed, when male mice were fed on LPD from P14 to P56, they were still stunted at P91 (Figure 2C), despite the removal of the nutritional stress. It is noteworthy that, despite the prevalence of stunting in male mice at P91, mice were not wasted after refeeding (Figure 2G-H). Notably, we did not observe any evidence of differential food intake between the experimental groups that could account for the observed phenotypes (Supplementary Figure 2A-B). However, caution is warranted when interpreting these results, as the data were obtained by extrapolating the amount of remaining food in each cage after one week.

**Figure 2:**
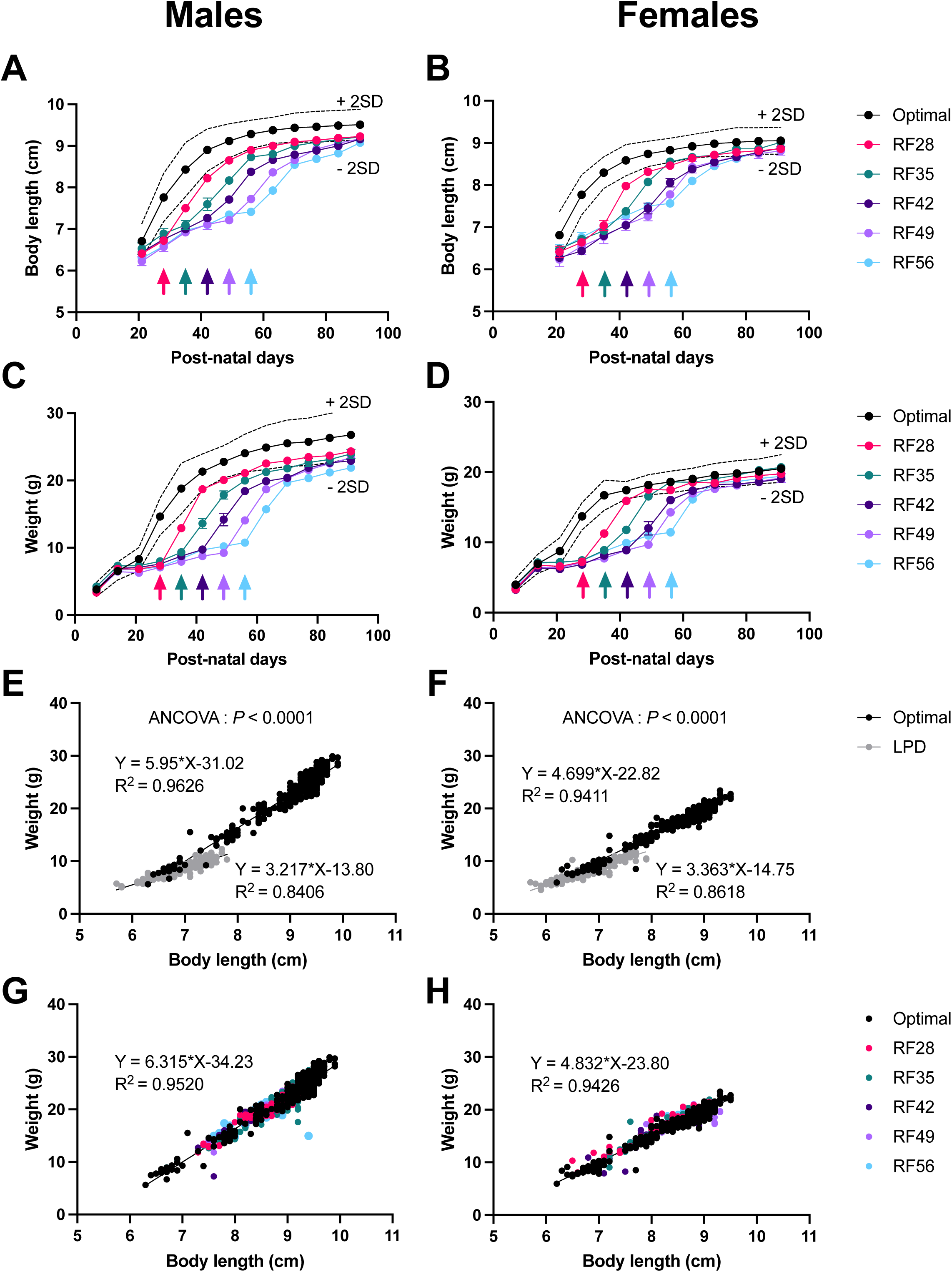
Chronically malnourished juvenile male and female mice differ in their dynamics of growth. Body length (**A, B**) and weight (**C, D**) of male and female optimal diet- or low-protein diet-fed mice refed with optimal diet respectively at post-natal day 28 (RF28), 35 (RF35), 42 (RF42), 49 (RF49), or 56 (RF56). Arrows correspond to the day of refeeding. **(E, F)** Length-versus-weight of optimal- and low-protein diet-(LPD) fed animals. (**G, H**) Length-versus-weight of optimal-fed and refed mice, after the time of refeeding. SD: standard deviation of optimal-fed group. Data are mean ± SEM. N = 8-10 mice were used in the refeeding groups. To ensure time-matched blood sampling in the optimal-fed group, each mouse was bled only three times, requiring N = 28 males and N = 35 females.

Using the weight-for-age (WA) relationship described by Anstead et al. (22), based on the classification by Gómez et al. (25), we calculated WA in all our groups (Figure 3A-B). After weaning, WA was inferior to 50% in LPD-fed male mice (Figure 3A), showing that the mice suffered from severe malnutrition. In females, however, LPD induced severe malnutrition, but the severity was corrected with time, reaching moderate malnutrition at P56 (Figure 3B). Refeeding with chow diet in male mice only partially corrected the severity of malnutrition, since RF42 mice had a WA score of 85.1 ± 2.1% (5% confidence interval), going as low as 81.6 ± 3.0% for RF56 mice, meaning they still faced mild malnutrition. As was the case previously, refeeding malnourished female mice completely corrected early-life malnutrition (Figure 3B). Overall, our data show a good example of nutritional imprinting of PEM, especially in males. Growth in mice occurs with a linear phase that lasts about two weeks after weaning followed by a plateau phase (26). We observed that this pattern was similar after refeeding. We thus compared growth catch-up by comparing the slopes following refeeding (or weaning, in the case of the optimal-fed group) (Figure 3C-D). In males, growth catch-up differed significantly between the groups (*P* = 0.0205; ANCOVA). After weaning on optimal diet, mice gained on average 0.11 cm.day^-1^. This value was similar when mice were refed after only one week on LPD (group RF28), but differed significantly with all other groups, which had a catch-up rate of about 0.08 cm.day^-1^ (Figure 3C). Thus, in males, growth catch-up after refeeding is impaired by long-term exposure to LPD in early life. In females, however, growth catch-up did not differ significantly between groups (*P* = 0.0504; ANCOVA; Figure 3D), showing a sex-dependent phenotype related to resilience to LPD feeding in early life.

**Figure 3:**
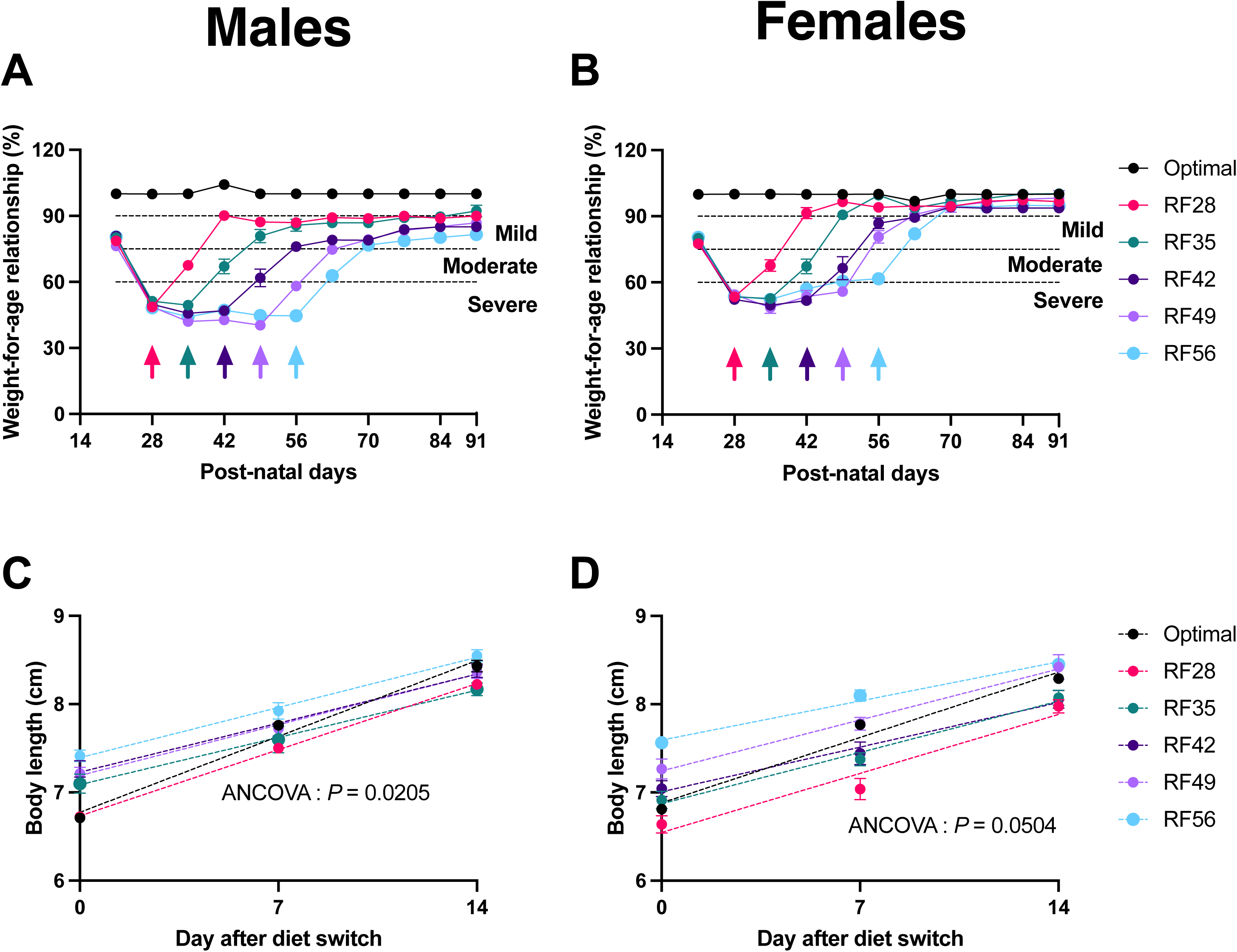
Optimal refeeding after long-term juvenile malnutrition is insufficient to fully recover systemic growth in males, but not in females. (A, **B)** Weight-for-age relationship of male and female optimal diet- or low-protein diet-fed mice refed with optimal diet respectively at post-natal day 28 (RF28), 35 (RF35), 42 (RF42), 49 (RF49), or 56 (RF56). Arrows correspond to the day of refeeding. **(C, D)** Body length evolution during the two weeks following refeeding, or from day 21 to day 35 in optimal-fed mice. Data are mean ± SEM. N = 8-10 mice were used in the refeeding groups. To ensure time-matched blood sampling in the optimal-fed group, each mouse was bled only three times, requiring N = 28 males and N = 35 females.

Furthermore, we measured weight of liver, gastrocnemius muscle and subcutaneous and perigonadal fat, and length of femur, tibia, small intestine and colon (Figure 4). Low-protein diet after weaning was associated to an increase of relative mass of subcutaneous fat (males: Figure 4B; females: Supplementary Figure 3B). In males, we observed significantly reduced femur and tibia size (Figure 4E-F), along with reductions in intestinal and colon sizes, with the severity of these reductions increasing with longer durations on the LPD (Figure 4G-H). When we plotted the totality of the measured parameters using Principal Component Analysis, we saw that values were mostly spread across PC1, which accounted for 30.44% of the total variance (Figure 4I). While optimal-fed animals clustered separately from refed mice, the latter formed a cluster, showing a biological difference between optimal-fed and refed mice. In contrast, in females, we only saw slight variations of fat deposits, but all points clustered together in a Principal Component Analysis (Supplementary Figure 3I).

**Figure 4:**
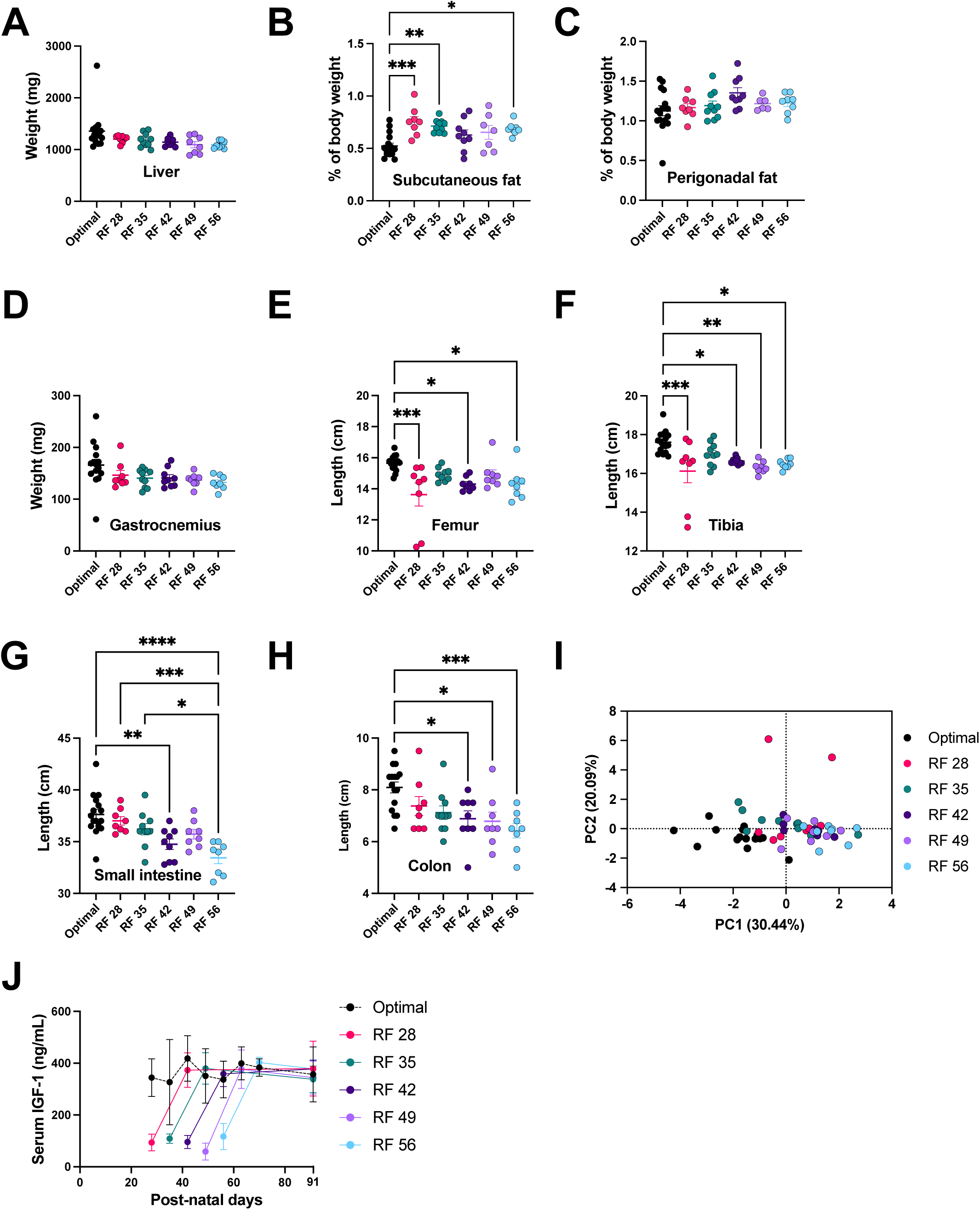
Optimal refeeding after early-life malnutrition in males is insufficient to fully recover organ growth. Weight of liver **(A)**, subcutaneous fat **(B)**, perigonadal fat **(C)**, and gastrocnemius muscle **(D)**; length of femur **(E)**, tibia **(F)**, small intestine **(G),** and colon **(H)** of male optimal diet- or low-protein diet-fed mice refed with optimal diet respectively at post-natal day 28 (RF28), 35 (RF35), 42 (RF42), 49 (RF49), or 56 (RF56). **(I)** Principal Component Analysis of parameters measured in **A** to **H**. **(J)** Serum concentration of IGF-1 measured at the switch of diet, 2 weeks after switch and at the end of the experiment in aforementioned mice. Data are mean ± SEM. Optimal: N = 16; refed: N = 8 to 10 mice. To ensure time-matched blood sampling in the optimal-fed group, N varies in panel **J**. Hence, for the optimal-fed group, N = 5 at P28 and P35; N = 14 at P42, P49 and P56; N = 6 at P63 and P70; N = 12 at P91. * *P* < 0.05; ** *P* < 0.01; *** *P* < 0.001; *** *P* < 0.0001, one-way ANOVA followed by Tukey’s post-hoc test.

Insulin-like Growth Factor 1 (IGF-1) is a hormone mainly produced by the liver as a downstream effector of growth hormone secretion (27). Circulating levels of IGF-1 directly regulate bone growth during juvenile age (28), and IGF-1 secretion is reduced upon chronic undernutrition (29). We thus measured circulating IGF-1 in males at three timepoints: at diet switch, two weeks after (at the end of the linear growth phase), and at P91 when the mice were killed (Figure 4J). We took time-matched samples in optimal-fed mice. In optimal-fed mice, IGF-1 levels did not differ significantly from P28 to P91 (about 350 ng.mL^-1^). In contrast, all mice fed LPD had an IGF-1 serum concentration around 100 ng.mL^-1^ at the time of refeeding, capturing an important signature of stunting in our model of PEM. Two weeks after refeeding, IGF-1 levels were comparable to those of optimal-fed male mice, and did not significantly vary at P91. Thus, while refeeding with an optimal diet stimulated IGF-1 secretion, it was not enough to compensate for the growth retardation, suggesting irreversible effects on male growth induced by long-term exposure to LPD after weaning.

### Lp^WJL^ treatment during the refeeding fails to improve systemic growth

Given the sex-related differences observed previously, where refeeding with an optimal diet fully compensated growth delay in females even after 56 days, we focused on male mice to explore the phenotype in a context where growth catch-up did not fully occur. We chose the most severe stunting phenotype that we observed after refeeding, i.e., mice that were refed at P56. We separated mice in three experimental groups: mice fed optimal diet from P14 to P119, mice fed LPD from P14 to P56, then optimal diet from P56 to P119, and mice fed LPD during the same period of time. We decided to keep until day 119, to see if the stunting phenotype observed at day 91 was still present four weeks later.

We previously reported that the Lp^WJL^ strain stimulates growth of juvenile mice upon chronic malnutrition (16,17). In these studies, treatment with Lp^WJL^ was performed in a model where, besides protein, fat was also drastically reduced in the diet. Treatment was performed after weaning, i.e., during the linear phase of growth. We wondered about the effect of Lp^WJL^ upon PEM and hypothesized that treating our mice during the refeeding phase, which corresponds to a linear phase of growth, would stimulate their growth to a point where mice would totally catch up their growth after a period of protein scarcity. As expected, mice fed LPD all the time showed severe stunting and underweight (Figure 5A-B). Despite a significant increase in body weight and length after refeeding, placebo-treated refed mice (RF56, Figure 5A-B) still presented signs of stunting and underweight, since their body length was less than 2 standard deviations from the mean of the optimal-fed group. Contrary to what we had hypothesized, supplementation with Lp^WJL^ during the refeeding phase did not have an additive effect on growth or weight, with only a slight increase in weight in LPD-fed mice after P91. As described in the previous section, we found no evidence of altered food intake in any of the experimental conditions (Supplementary Figure 2C).

**Figure 5:**
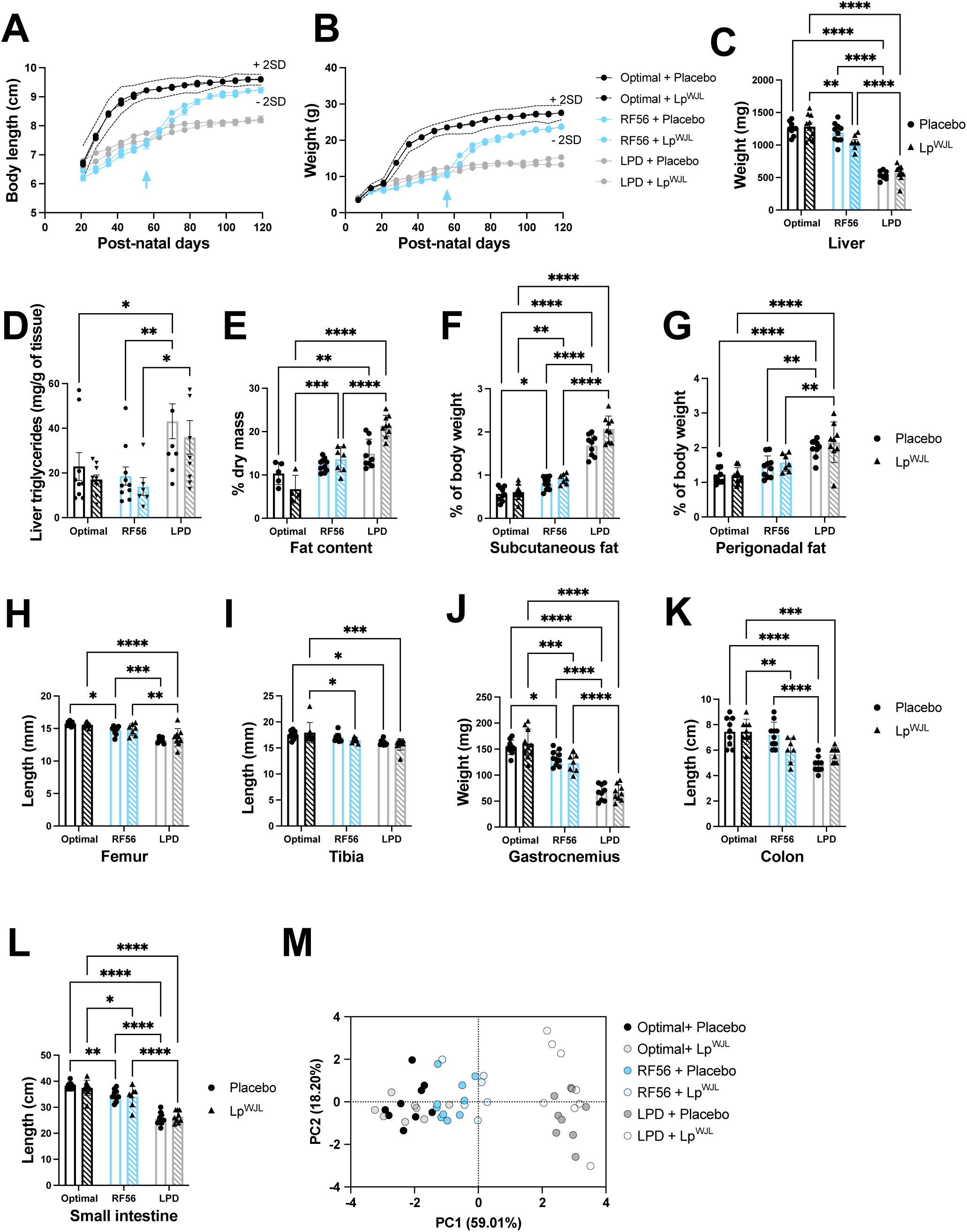
Lp^WJL^ treatment during the refeeding phase fails to improve systemic growth. Body length **(A)** and weight **(B)**; weight of liver **(C)**; liver triglycerides **(D)**; fat content **(E)**; weight of subcutaneous fat **(F)** and perigonadal fat **(G)**; length of femur **(H)**, tibia **(I)**, and weight of gastrocnemius muscle **(J)**; length of colon **(K)**, and small intestine **(L)** of male mice refed at post-natal day 56 (RF56) or maintained on low-protein diet (LPD) and treated with *Lactiplantibacillus plantarum* WJL (Lp^WJL^) or a placebo. **(M)** Principal Component Analysis of parameters measured in **C** and **F** to **L**. Data are mean ± SEM. N = 7-10 mice per experimental group. * *P* < 0.05; ** *P* < 0.01; *** *P* < 0.001; *** *P* < 0.0001, two-way ANOVA followed by Šídák’s post-hoc test.

As we show in Figure 5D, refed mice exhibited liver triglyceride (TG) levels similar to those of mice fed the optimal diet, while the low-protein diet (LPD) resulted in increased liver fat accumulation. Additionally, no significant effect on liver TG levels was observed with Lp^WJL^ treatment. To our surprise, Lp^WJL^ treatment had no impact on size and weight of several organs (Figure 5E-L). This was evident when running Principal Component Analysis (Figure 5M), where mice clustered mainly along PC1 (59.01% of total variance) according to diet, but not to treatment. Thus, treatment with Lp^WJL^ during the refeeding phase failed to further stimulate growth after prolonged exposition to protein scarcity, highlighting the long-lasting deleterious effects of protein malnutrition during early life.

### Suboptimal refeeding with a non-obesogenic Western diet compensates early-life associated stunting, but impairs glucose metabolism

One of the main drivers of malnutrition is suboptimal nutrition, defined as inadequate nutrient intake that does not meet the age-specific recommendations (30). Finally, we sought to characterize the impact on weight gain and growth of LPD feeding followed by refeeding with a suboptimal diet. For this experience, we decided to refeed the animals at P35, i.e., the first time when growth catch-up was impaired upon refeeding with an optimal diet (Figure 2A-B). Indeed, we sought to establish a novel model of “double burden of malnutrition”, defined as the simultaneous manifestation of both undernutrition and overweight (31). During the first burden (P14 to P35), mice were fed optimal diet or LPD. During the second burden (P35 to P119), mice were fed two types of suboptimal diet: Total New Western Diet (WD) or a modified version of WD (MWD), with protein accounting for 7.5% of total kcal (instead of 15.3%). In parallel to these groups, we also fed mice optimal diet, WD, or MWD from P14 to P112 (Figure 1C). Unlike diet-induced obesity diet traditionally used in nutritional studies, WD has a variety of fat sources and is more similar to the human Western diet. It is noteworthy that this version of WD is not obesogenic in C57Bl/6J mice (21), so the idea was to provide a suboptimal diet without strongly affecting body weight. The mice shown in Figures 6 and 7 received either placebo or Lp^WJL^, as detailed in the Methods section. For clarity, Figure 6 presents data from placebo-treated mice only, while Figure 7 includes data from both placebo- and Lp^WJL^-treated mice. The placebo-treated mice in Figure 6 are the same as those in Figure 7.

**Figure 6:**
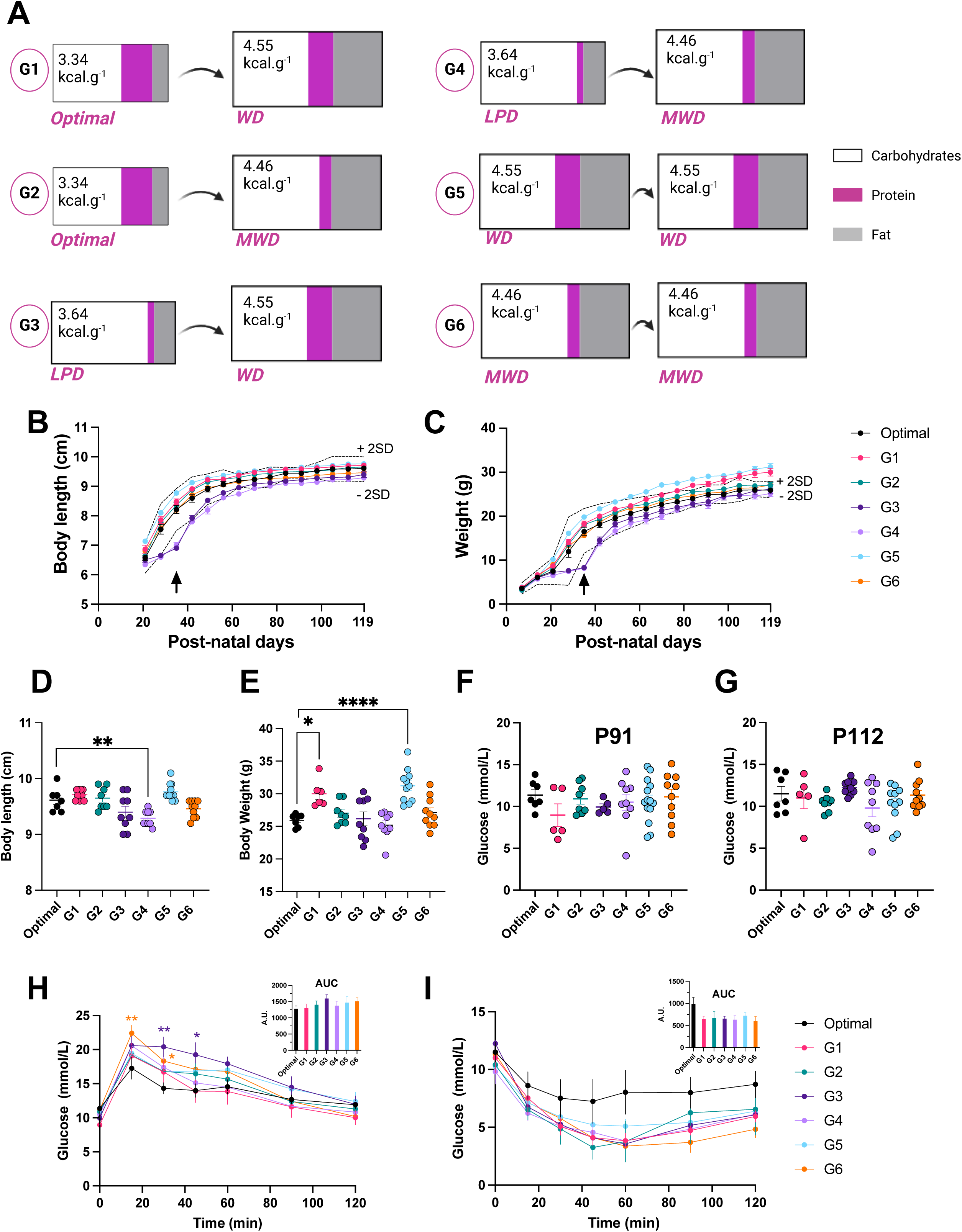
Suboptimal refeeding with a non-obesogenic Western diet compensates early life-associated stunting, but impairs glucose metabolism. **(A)** Experimental groups used in the procedure, with the respective diets. Body length **(B)** and weight **(C)** of mice. The arrow corresponds to the time of diet shift. Length **(D)** and weight **(E)** at post-natal day 119. Plasma glucose after 6 hours of food deprivation at post-natal day 91 **(F)** and 112 **(G)** of mice described in panel **A**. Intraperitoneal glucose tolerance test **(H)** and insulin tolerance test **(I)** performed at post-natal day 91 and 112 respectively on mice described in panel **A**. Inset: area under the curve (AUC) of the respective tests. Data are mean ± SEM. N = 7-10 mice per experimental group. * *P* < 0.05; ** *P* < 0.01; *** *P* < 0.0001, one-way ANOVA followed by Dunnett’s post-hoc test vs. optimal-fed mice.

**Figure 7:** Supplementation with Lp^WJL^ in suboptimal conditions further impairs glucose tolerance after early-life protein restriction. Intraperitoneal glucose tolerance test (GTT) and insulin tolerance test (ITT) performed respectively at P91 and P112 on male mice treated with *Lactiplantibacillus plantarum* WJL (Lp^WJL^) or a placebo from P35 to P119. Experimental groups are described in Figure 6A. Placebo-treated mice are the same as those presented in Figure 6. Data are mean ± SEM. N = 9-10 mice per experimental group. * *P* < 0.05; ** *P* < 0.01, Student’s *t*-test with Welch’s correction.

During the first burden, mice on LPD gained significantly less weight and were stunted (Figure 6B-C; G3: mice switching from LPD to WD, and G4: mice switching from LPD to MWD). However, upon refeeding with WD or MWD, mice quickly gained weight and had a significant growth catch-up (Figure 6B-C; days 35 to 119). Indeed, at P119 all groups of mice had a similar body length (Figure 6D). Feeding with WD after optimal diet induced a significant gain in body weight compared to optimal-fed animals (Figure 6E), but this was not the case after LPD. In contrast, MWD did not induce a significant increase in body weight, even when mice were fed MWD from P14 onwards. It is noteworthy that weight gain did not correlate with increased food intake, as mice in the group with the highest caloric intake (G4) had weights similar to those of optimally fed mice. Conversely, the overweight mice in groups G1 and G5 showed no evidence of elevated food intake (Supplementary Figure 2D).

To assess whether LPD followed by WD or MWD dysregulates glucose homeostasis, we evaluated glucose and insulin tolerance. At P91 or P112 respectively, after 6 hours of food deprivation, no diet had a significant impact on glycemia (Figure 6F-G). It is noteworthy that glycemia did not differ significantly between the two time points (two-way ANOVA: effect of time, *P* = 0.81). When we performed an intraperitoneal glucose tolerance test (GTT), we saw that, unlike WD, MWD feeding after from P14 onwards (group 6), induced a significant glycemic increase in the early phase of the GTT (the so-called glycemic excursion) (Figure 6H, G6). However, this was not the case when MWD was given after P35 (Figure 6H, G2 and G4). Interestingly, although WD alone or following chow feeding did not alter glucose tolerance (Figure 6H, G1 and G5), early-life priming with LPD led to a temporary glucose intolerance (Figure 6H, G3). As a whole, LPD-fed mice were susceptible to develop early-phase glucose intolerance when they were refed with WD, i.e., a suboptimal diet rich in fat.

To infer if this phenotype was due to a change in insulin sensitivity, we performed an insulin tolerance test (ITT). We saw that insulin tolerance was not modified by any of the different diet combinations (Figure 6I), highlighting that the aforementioned glucose intolerance was not due to impaired insulin sensitivity. Overall, we show that, while it does not affect growth catch-up after protein malnutrition-mediated early life stunting, suboptimal nutrition leads to impaired glucose control later in life.

### Supplementation with Lp^WJL^ in suboptimal nutritional conditions further impairs glucose tolerance after early-life protein restriction

In the same line as previously, we investigated the effect of a supplementation with Lp^WJL^ during the second burden of the aforementioned protocol (i.e., from P35 to P119). We found no effect of adding Lp^WJL^ on weight gain and systemic growth (Supplementary Figure 4). Likewise, two-way ANOVA analysis did not reveal any significant effect of Lp^WJL^ on organ size or weight (Supplementary Figure 5). We further assessed glucose metabolism by performing a GTT and an ITT. We previously showed that Lp^WJL^ treatment resulted in improved glucose tolerance during the post-weaning period in chronic malnutrition mice (17). It is noteworthy that, in our experimental setting, treatment with Lp^WJL^ did not affect glycemia after a 6-hour-long food deprivation (Supplementary Figure 6; effect of Lp^WJL^, P = 0.0869, two-way ANOVA).

Overall, glucose tolerance was not affected in mice fed optimal diet after weaning (G1 and G2 mice, Figure 7A-D), nor mice that were fed WD or MWD all long (G5 and G6 mice; Figure 7I-L). Surprisingly, mice facing a double burden of malnutrition, particularly those fed WD after LPD (G3 mice; Figure 7E-F), showed significant glucose intolerance when given Lp^WJL^ during the second burden. Furthermore, Lp^WJL^ did not significantly affect insulin tolerance in any of the groups (Figure 7). Overall, these findings indicate that Lp^WJL^ treatment exacerbates glucose intolerance in a mouse model of a double burden of malnutrition manifested by early protein malnutrition followed by western diet nutrition.

## DISCUSSION

Early-life PEM has fatal consequences and is one of the leading causes of infant mortality in the developing world (30). Maintenance of adequate nutrition and rehabilitation are the most effective interventions to reduce the deleterious effects of PEM (32). Undernutrition in infants during the first 1,000 days is influenced by infections, subclinical pathogen carriage, and altered gut microbiota. This period is critical for healthy growth and development (12). In our mouse model, we defined P35 as the equivalent of this critical period in humans, while Experiment 1 extended malnutrition to P56 to study its long-term effects. In this study, we aimed at establishing a mouse model of PEM, followed by a nutritional rehabilitation, in order to explore the long-lasting consequences of early-life PEM.

Early-life PEM results in stunting (33). In our protein malnutrition model, slow growth and weight gain rate occurred after weaning, with over 50% reduction after 5 weeks on LPD. In humans, the two main clinical syndromes of the extreme forms of children with PEM are marasmus and kwashiorkor (34). While marasmus results from adaptation to severe depletion of calories and nutrients, kwashiorkor (“sickness of the weaning” in Ga language) is the consequence of a diet with inadequate protein but reasonably normal caloric intake. Although our mouse model is characterized by a diet low in protein but not in calories, it does not replicate either form of human juvenile malnutrition, as it lacks hallmark features such as edema and hepatomegaly (34), probably due to the fact that our mice are not in contact with any enteric pathogens and thus are not susceptible to infection and inflammation. Indeed, several studies have shown that malnourished mice are more susceptible to infection, thus mimicking one of the hallmarks of human malnutrition (22,23). Furthermore, nutritional rehabilitation after kwashiorkor results in increased IGF-1 levels, but the values are lower than in healthy children (35). In our mouse model, IGF-1 levels are fully restored by nutritional rehabilitation, suggesting that mice are more resilient than humans to early-life nutritional stress. This is confirmed by the fact that, in our model, mice are kept under severe protein malnutrition for several weeks after weaning. While this results in a severe phenotype regarding growth and weight, our mice did not show any particular sign of torpor, reduced activity, or change in appetite during the nutritional challenge. Moreover, unlike Ferreira-Paes et al. (23), who describe a model of marasmus using weanling BALB/c mice, nutritional rehabilitation in male mice did not fully resolve the severity of protein malnutrition. Indeed, in our experimental setting, male mice were stunted after nutritional rehabilitation, although early-life wasting was corrected. It is noteworthy that the earlier the rehabilitation took place, the better the response was. Nevertheless, even after post-natal day 56 (i.e., young adult mice), growth occurred at a rate of 0.08 cm.day^-1^ after refeeding on an optimal diet, showing the remarkable capacity of mice to adapt to deleterious nutritional environments.

Recent advances in the knowledge of malnutrition have shown that the gut microbiota is a contributor to the deleterious stunting and wasting phenotype (36). Indeed, colonization of germ-free mice with kwashiorkor-associated microbiota in combination with a Malawian diet leads to severe weight loss (37). In our previous studies, we demonstrated that intestinal microbiota plays a critical role in postnatal growth, and specifically, that treatment with Lp^WJL^ from P21 to P56 had beneficial effects on growth through modulation of IGF-1 and insulin production (16,17). In the present study, however, Lp^WJL^ was administered later, from P56 to P119, during the recovery phase from protein undernutrition. In the current model, IGF-1 levels are fully restored by nutritional rehabilitation alone, suggesting that mice are more resilient to early-life nutritional stress compared to humans. This difference suggests a potential age-dependent window during which microbiota interventions, such as Lp^WJL^ treatment, may exert the strongest influence on IGF-1 and insulin pathways, and underscores the complexity of microbial interactions across different stages of development.

Overall, our results confirm the Developmental Origins of Health and Disease (DOHaD) paradigm, which stipulates that developmental and early postnatal insults, such as malnutrition, can have deleterious impacts in adult life (38). In line with that, the DOHaD paradigm is particularly appropriate in understanding the nutritional transitions in countries that face the double burden of malnutrition. Therefore, we sought to establish a model with suboptimal refeeding after a moderately long period of low-protein diet. It is well known that severe acute malnutrition affects glucose homeostasis in infants (4,39), a feature that persists later in life (40,41). Furthermore, studies in rodents have shown that post-weaning PEM induces hepatic alterations, such as steatosis and impaired hepatic mitochondrial turnover (42–44). Interestingly, two independent studies using a high-fat diet (58% kcal from fat) after LPD have shown that PEM after weaning does not potentiate fat accumulation, hepatic steatosis and insulin resistance in adult young mice (42,45). In their mouse model of double burden of malnutrition, Dalvi et al. described that LPD feeding for 4 weeks post-weaning induces transient glucose intolerance after a recovery on chow feeding (42), showing that LPD priming induces deleterious effects later in life associated to poor metabolic health. Indeed, in our model, while we saw no differences in growth with mice on MWD after LPD, LPD priming also induces deficiencies in glucose control later in life. Although results on mouse models are still inconsistent and fail to fully recapitulate the metabolic alterations observed in malnourished humans, they provide key mechanistic insights into the different pathologies associated to stunting in early life.

Indeed, cohorts of malnourished mice and humans have shown that growth and metabolic functions are impaired, associated to alterations in the gut microbiota, and that this impaired phenotype could be transferred (36). Similarly, extensive research has shown that the overnourished microbiota is associated to loss of richness of taxa and enrichment of bacteria associated to inflammation (46,47). While most studies have been correlative on this issue, they have given an insightful look into the alterations of microbial ecology in cases of malnutrition. Little is known on humans facing the double burden of malnutrition. A 2018 study on a Mexican cohort of 36 patients (undernourished, obese or controls) showed similarities in the composition of the microbiota between the obese and undernourished group. In line with these findings, we hypothesized that treatment with Lp^WJL^ could have a beneficial impact in mice facing a double burden of malnutrition. Not only did we not see any beneficial effect of Lp^WJL^ in this context, but the treatment induced further impairment of glucose tolerance in mice fed a suboptimal Western diet. Thus, caution should be used when developing probiotic strategies for PEM when infants also face a double burden of malnutrition.

Overall, we provide a new model of refeeding after early-life PEM, where we show sex-associated differences in the long-term response to refeeding. We provide data showing that suboptimal refeeding with a moderately low-protein diet, which is also rich in fat, is sufficient to induce growth in malnourished mice, although this is associated to metabolic perturbations. Further investigations into the specific mechanisms driving these features will help uncover mechanisms associated to PEM and the double burden of malnutrition.

## DATA AVAILABILITY

Data can be provided upon request.

## ACKNOWLEDGEMENTS

This work was supported by funding from Fondation pour la Recherche Médicale (Équipe FRM EQU202203014629). FDV was a recipient of a Young Researcher Prize from Fondation des Treilles. The authors declare no conflict of interest associated to this study.

JLT, FL, and FDV designed research; JLT, AC and DS conducted research; JLT and FDV analyzed data; and JLT, FL and FDV wrote the paper. FDV had primary responsibility for final content. All authors read and approved the final manuscript.

## List of abbreviations

GTT: Glucose tolerance test
IGF-1: Insulin-like growth factor 1
ITT: Insulin tolerance test
Lp^WJL^: *Lactiplantibacillus plantarum*
WJL LPD: Low-protein diet
PC: Principal component
PEM: Protein energy malnutrition
MWD: Modified Western Diet
WA: weight-for-age relationship
WD: Total New Western Diet

---

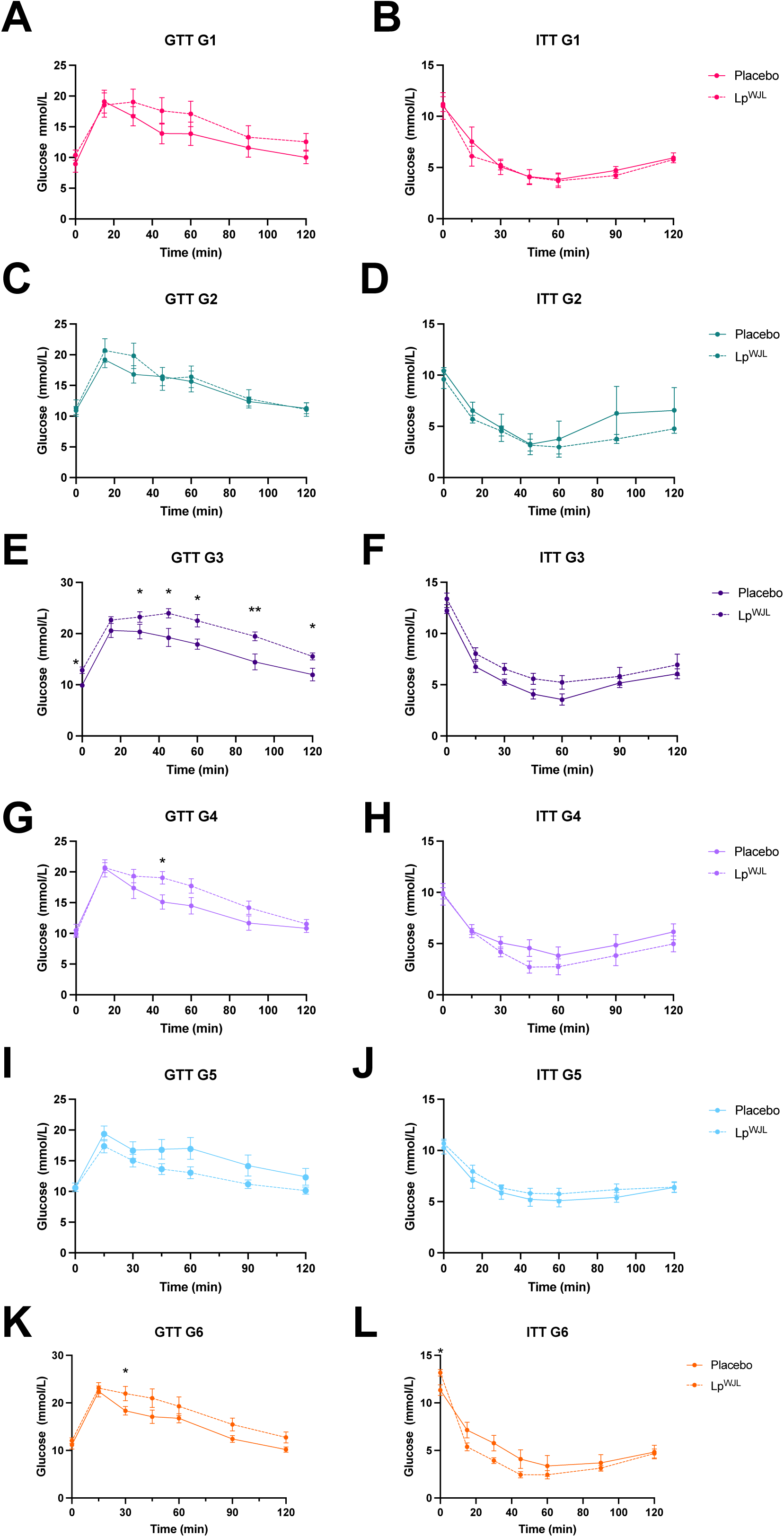

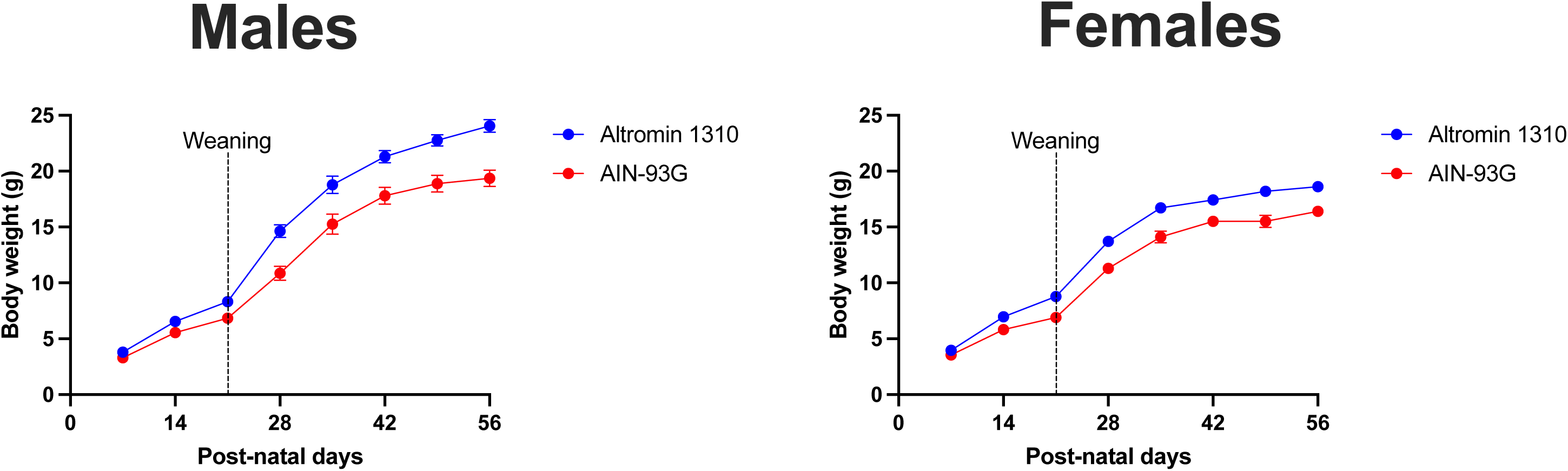

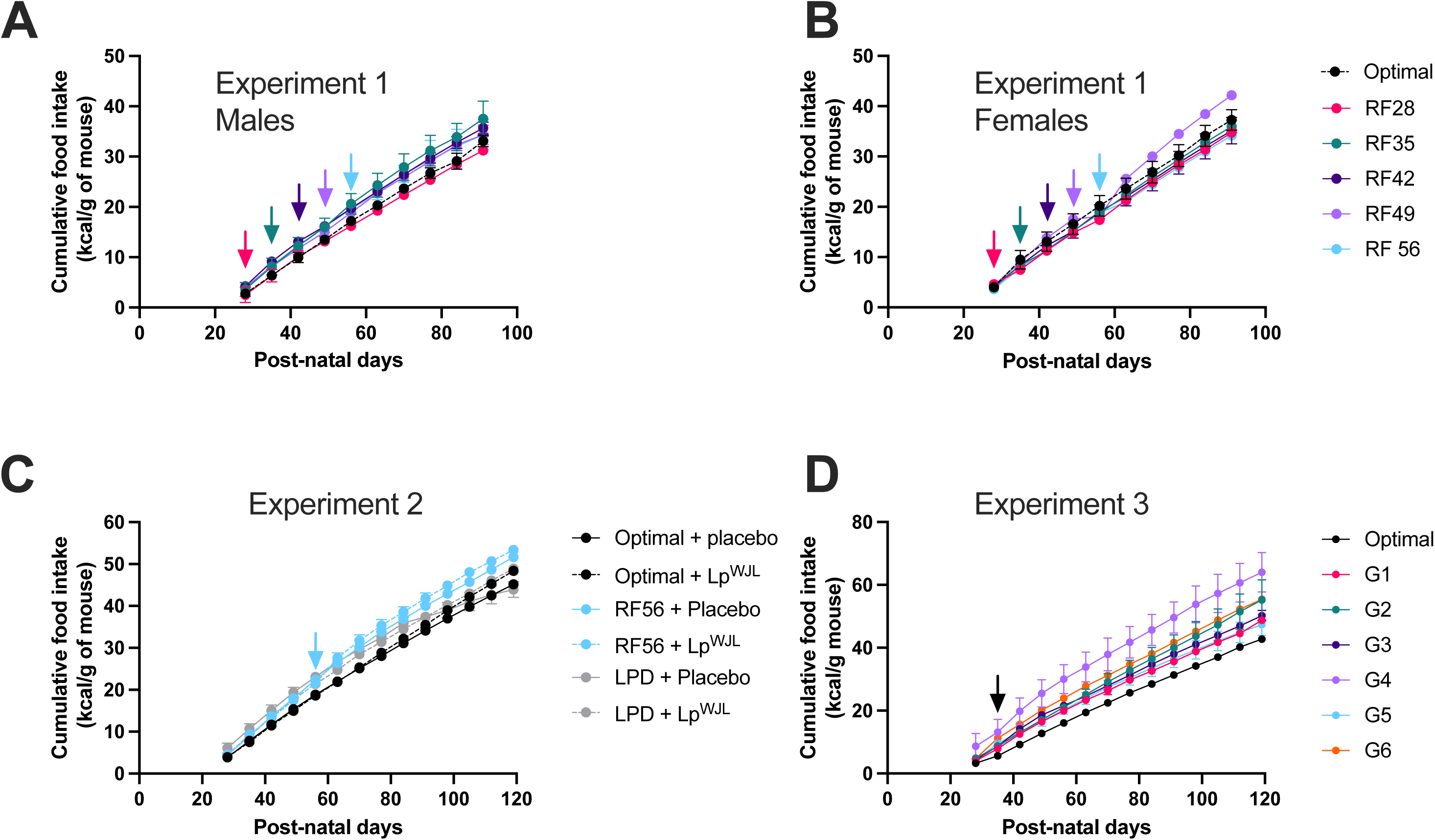

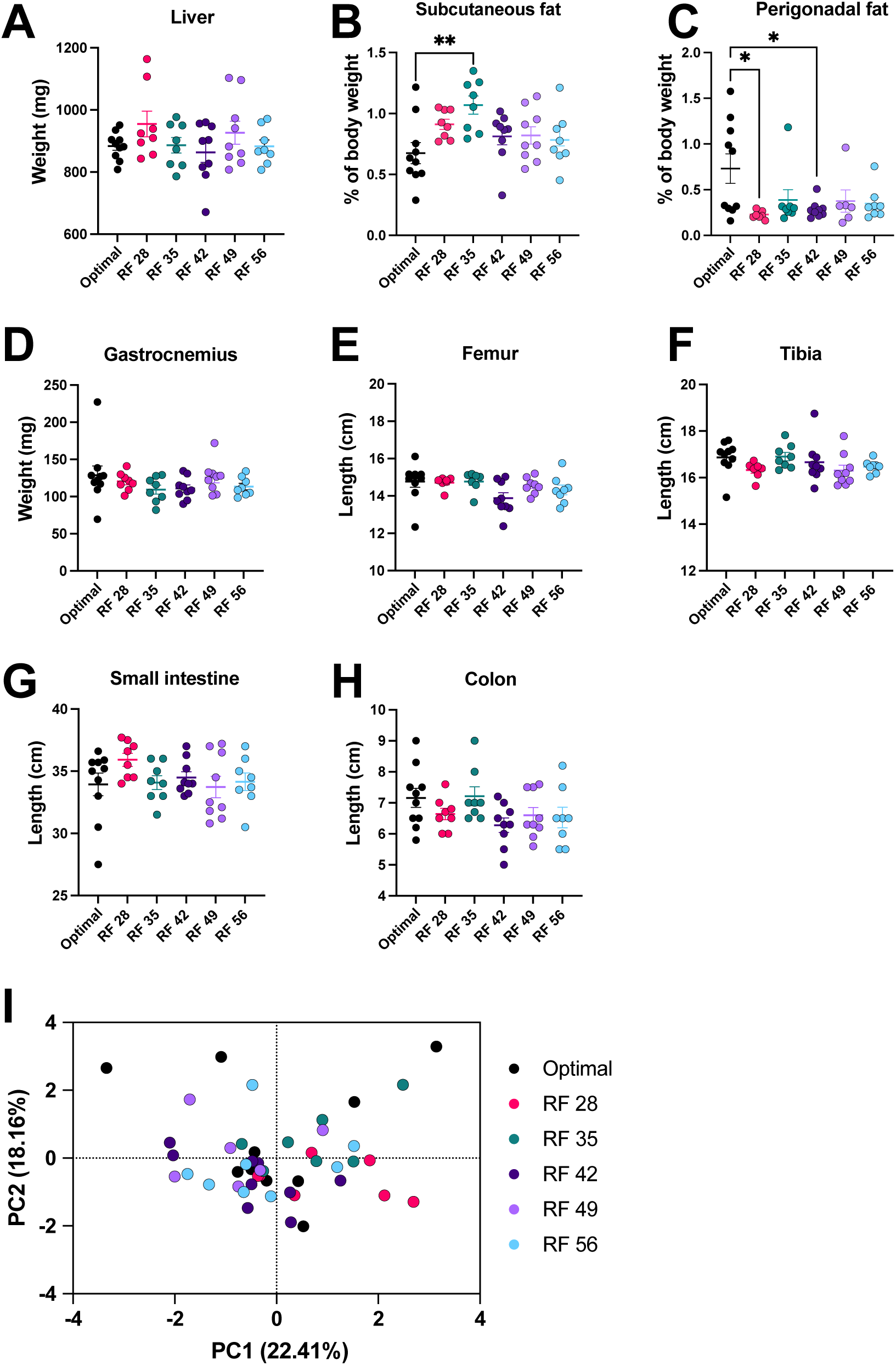

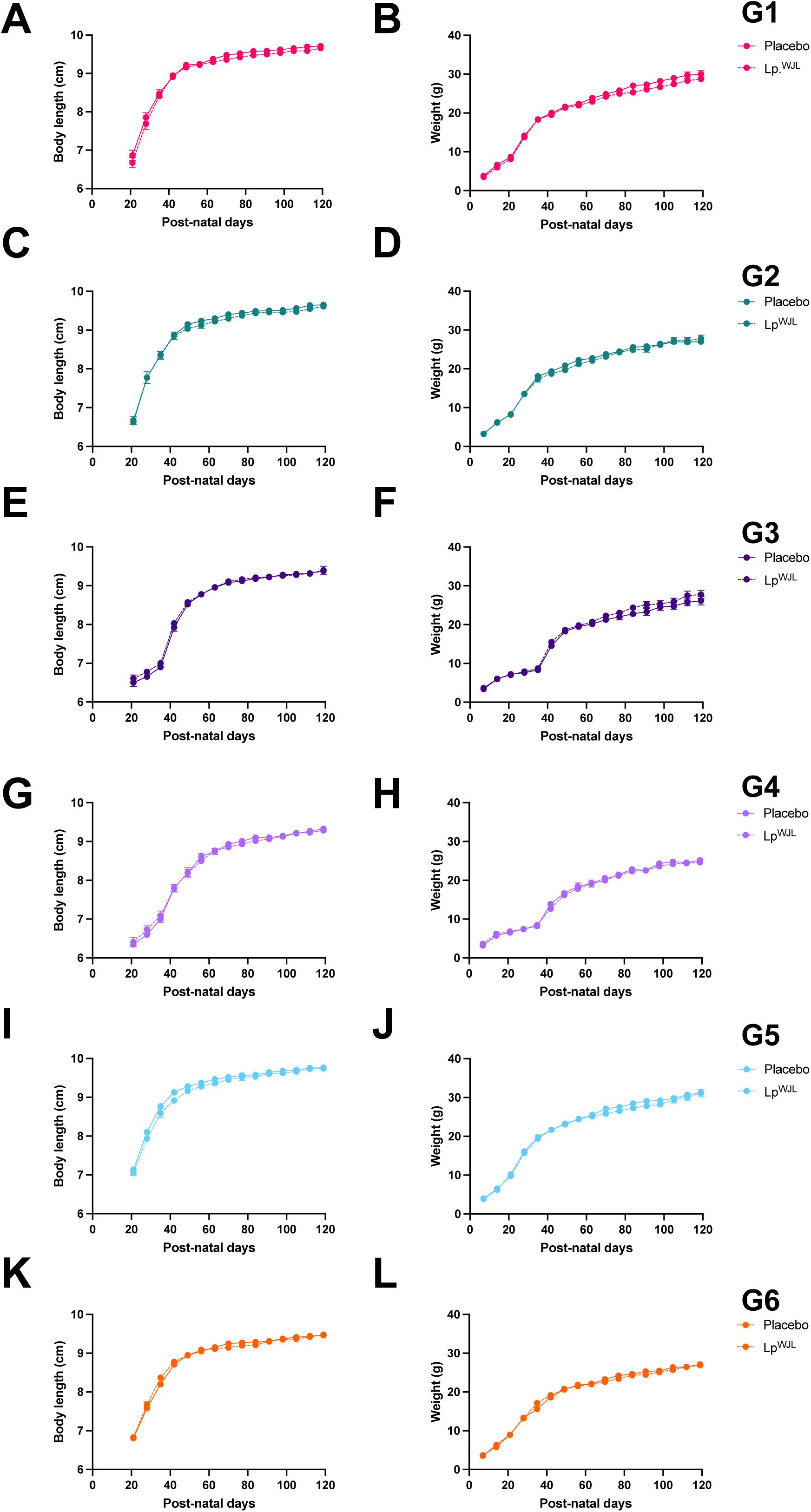

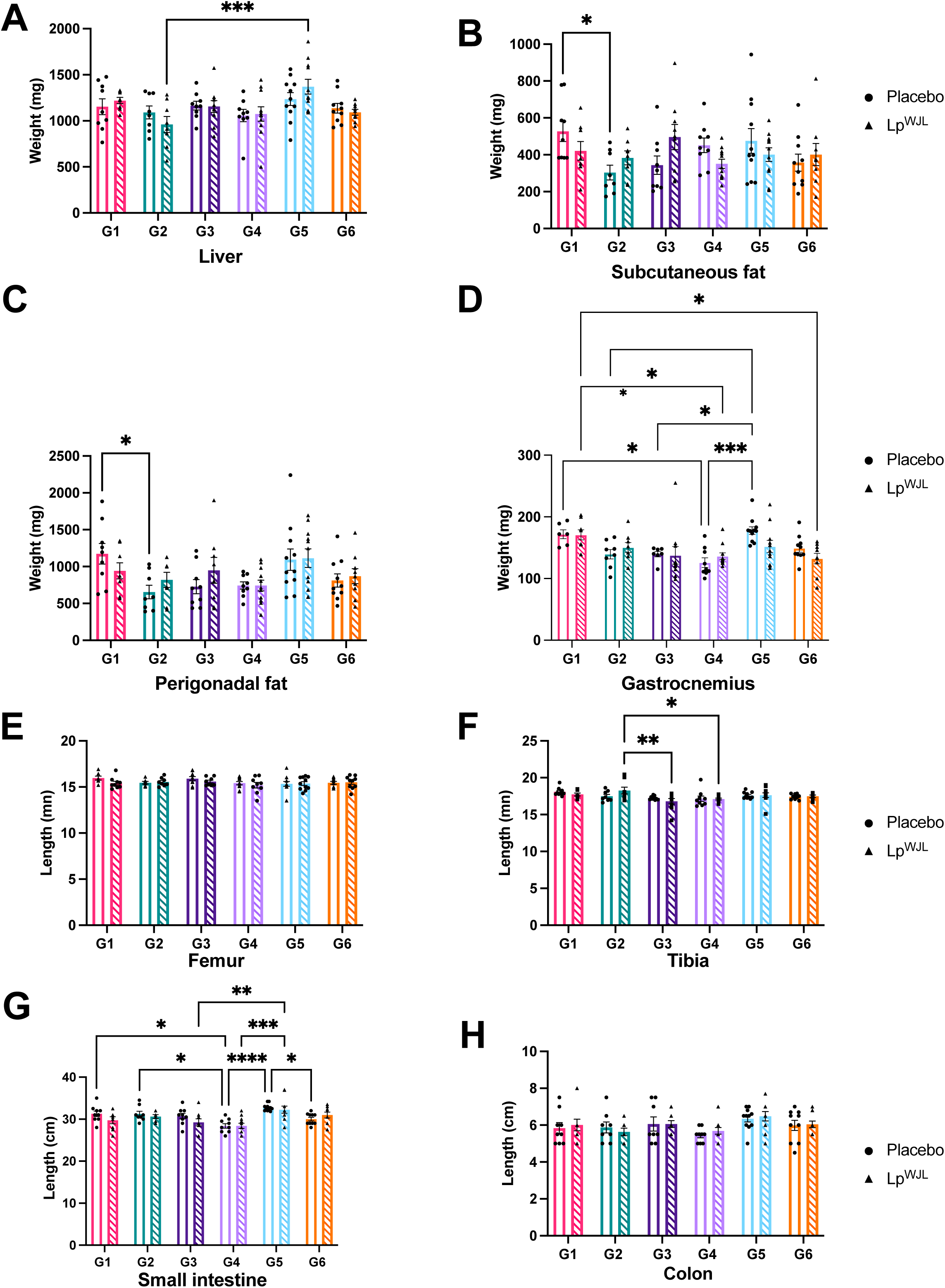

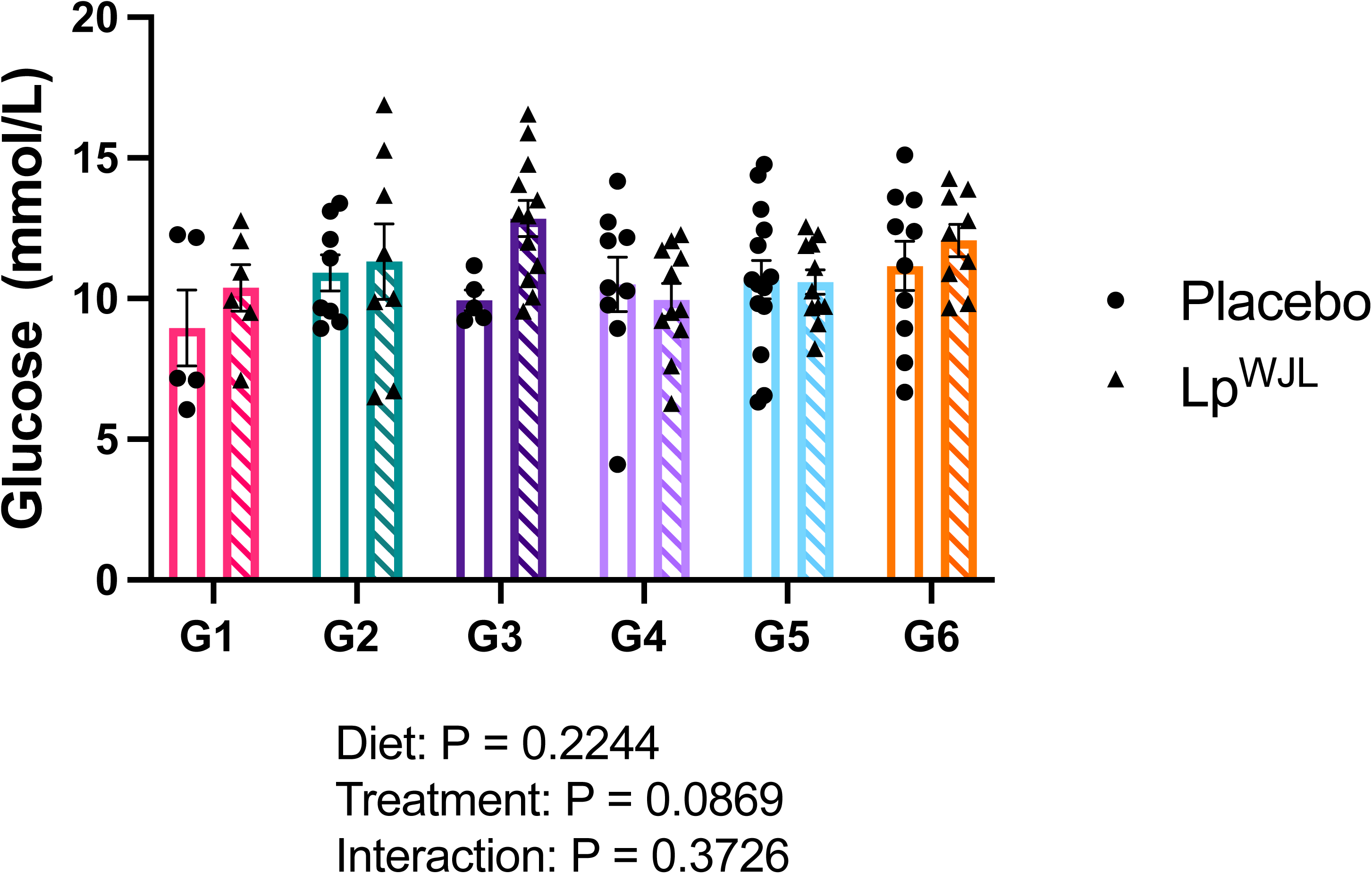

