## Supplementary Figure Legends for "Suboptimal refeeding compensates stunting in a mouse model of juvenile malnutrition"

**Supplementary Figure 1: Comparison of growth curves in C57Bl/6J mice fed AIN-93G or Altromin 1310.**

Growth curves for male (**A**) and female (**B**) mice show that those fed Altromin 1310 exhibit superior growth compared to those fed AIN-93G.

Data are mean ± SEM.

**Supplementary Figure 2: Food intake of mice in the different experimental settings.**

Cumulative food intake was measured in every cage by measuring the initial amount of food provided and the remaining food after one week, then calculating the average intake per mouse in each cage. Data are given for mice related to Figures 2-4 **(A)**, Figure 5 **(B)** and Figure 6 **(C)**. Arrows indicate dietary switch.

Data are mean ± SEM. N = 2-3 cages per group.

**Supplementary Figure 3: Optimal refeeding after early-life malnutrition in females compensates organ growth.**

Weight of liver **(A)**, subcutaneous fat **(B)**, perigonadal fat **(C)**, and gastrocnemius muscle **(D)**; length of femur **(E)**, tibia **(F)**, small intestine **(G),** and colon **(H)** of female control diet- or low-protein diet-fed mice refed with control diet respectively at post-natal day 28 (RF28), 35 (RF35), 42 (RF42), 49 (RF49), or 56 (RF56). **(I)** Principal Component Analysis of parameters measured in **A** to **H**.

Data are mean ± SEM. N = 8-10 mice per experimental group.

* *P* < 0.05; ** *P* < 0.01, one-way ANOVA followed by Tukey’s post-hoc test.

**Supplementary Figure 4: Supplementation with Lp^WJL^ during suboptimal refeeding does not impact systemic growth.**

Body length and weight of male mice treated with *Lactiplantibacillus plantarum* WJL *(*Lp^WJL^) or a placebo. Experimental groups are described in Figure 6**A**.

Data are mean ± SEM. N = 9-10 mice per experimental group.

**Supplementary Figure 5: Supplementation with Lp^WJL^ during suboptimal refeeding does not impact organ growth.**

Weight of liver **(A)**, subcutaneous fat **(B)**, perigonadal fat **(C)**, and gastrocnemius muscle **(D)**; and length of femur **(E)**, tibia **(F)**, small intestine **(G)** and colon **(H)** of male mice treated with *Lactiplantibacillus plantarum* WJL *(*Lp^WJL^) or a placebo. Experimental groups are described in Figure 6**A**.

Data are mean ± SEM. N = 9-10 mice per experimental group.

* *P* < 0.05; ** *P* < 0.01; *** *P* < 0.001; *** *P* < 0.0001, two-way ANOVA followed by Šídák’s post-hoc test.

**Supplementary Figure 6: Supplementation with Lp^WJL^ during suboptimal refeeding does not impact glycemia.**

Glycemia was measured at P91 after a 6-hour food deprivation in male mice treated with *Lactiplantibacillus plantarum* WJL *(*Lp^WJL^) or a placebo. Experimental groups are described in Figure 6**A**.

Data are mean ± SEM. N = 9-10 mice per experimental group. P-values are given after two-way ANOVA.
