## Supplementary material for "Suboptimal refeeding compensates stunting in a mouse model of juvenile malnutrition": Table 1

|  |  | Optimal diet (Altromin 1310) |  | Low-protein diet | Total New Western Diet (Teklad TD.140203) | Low-protein Total New Western Diet |
| --- | --- | --- | --- | --- | --- | --- |
| **Carbohydrates (g.kg^-1^)** | Monosaccharides | 0 | Corn starch | 405.786 | 232.9 | 284.022 |
|  | Disaccharides | 54.151 | Maltodextrin | 200 | 70 | 80 |
|  | Polysaccharides | 350.3 | Sucrose | 100 | 238.647 | 286 |
|  | Fiber | 45.48 | Cellulose | 130 | 30 | 50 |
|  | *kcal/g* | *1.98* |  | *2.82* | *2.2* | *2.5* |
|  | *kcal (% of total)* | *59.0* |  | *77.6* | *50.5* | *58.0* |
| **Protein (g.kg^-1^)** | Crude Protein | 225.155 | Casein | 46 | 190 | 92 |
|  |  |  | L-cystine | 0.7 | 2.85 | 1.4 |
|  | *kcal/g* | *0.90* |  | *0.19* | *0.7* | *0.3* |
|  | *kcal (% of total)* | *27.0* |  | *5.1* | *15.3* | *7.5* |
| **Fat (g.kg^-1^)** | Crude fat | 51.398 | Soybean oil | 70 | 31.4 | 31.4 |
|  |  |  | Anhydrous milk fat | 0 | 61.1 | 61.1 |
|  |  |  | Olive oil | 0 | 28 | 28 |
|  |  |  | Lard | 0 | 28 | 28 |
|  |  |  | Corn oil | 0 | 16.5 | 16.5 |
|  |  |  | Cholesterol | 0 | 0.4 | 0.4 |
|  | *kcal/g* | *0.46* |  | *0.63* | *1.5* | *1.5* |
|  | *kcal (% of total)* | *14.0* |  | *17.3* | *34.2* | *34.5* |
| **Energy density (kcal.g^-1^)** | | **3.34** |  | **3.64** | **4.4** | **4.3** |
|  | |  | Mineral mix (AIN93G-MX) | 35 | 13.4* | 13.4* |
|  | |  | Vitamin mix (AIN93G-VX) | 10 | 10 | 10 |
|  | |  | Choline bitartrate | 2.5 | 2.5 | 2.5 |
|  | |  | TBHQ, antioxidant | 0.014 | 0.028 | 0.028 |
|  | |  | Calcium phosphate, dibasic |  | 7.5 | 10.5 |
|  | |  | Calcium carbonate |  | 4.5 | 6.875 |
|  | |  |  |  | * Without Ca & P | |

**Table 1.** Composition of the different diets used in the study
